# PsiNorm: a scalable normalization for single-cell RNA-seq data

**DOI:** 10.1101/2021.04.07.438822

**Authors:** Matteo Borella, Graziano Martello, Davide Risso, Chiara Romualdi

## Abstract

Single-cell RNA sequencing (scRNA-seq) enables transcriptome-wide gene expression measurements at single-cell resolution providing a comprehensive view of the compositions and dynamics of tissue and organism development. The evolution of scRNA-seq protocols has led to a dramatic increase of cells throughput, exacerbating many of the computational and statistical issues that previously arose for bulk sequencing. In particular, with scRNA-seq data all the analyses steps, including normalization, have become computationally intensive, both in terms of memory usage and computational time. In this perspective, new accurate methods able to scale efficiently are desirable.

Here we propose *PsiNorm*, a between-sample normalization method based on the power-law Pareto distribution parameter estimate. Here we show that the Pareto distribution well resembles scRNA-seq data, independently of sequencing depths and technology. Motivated by this result, we implement *PsiNorm*, a simple and highly scalable normalization method. We benchmark *PsiNorm* with other seven methods in terms of cluster identification, concordance and computational resources required. We demonstrate that *PsiNorm* is among the top performing methods showing a good trade-off between accuracy and scalability. Moreover *PsiNorm* does not need a reference, a characteristic that makes it useful in supervised classification settings, in which new out-of-sample data need to be normalized.

*PsiNorm* is available as an R package available at https://github.com/MatteoBlla/PsiNorm

## 1 Introduction

Gene expression data exhibit a scale-free power-law distribution (*k*^*−λ*^) with the exponent fluctuating from 1 to 3. This result holds independently of experimental techniques (such as SAGE, microarray and RNA-seq experiments) and across different organisms (Nacher and Akutsu, 2006; Furusawa and Kaneko, 2003; Awazu *et al*., 2018; Ueda *et al*., 2004; Kuznetsov *et al*., 2002).

A power-law distribution has the property that large numbers are rare, while smaller numbers are more common. In transcriptomics this translates to the presence of a relatively low number of genes with high expression levels along with many low-abundant genes. This suggests the presence of a complex organization conserved among species (Barabási and Albert, 1999).

Supported by this observation, Lu *et al.* (2005) and Wang (2020) proposed two normalization methods based on Zipf’s law, a type of power law, for microarray and RNA-seq data, respectively, showing promising results. Zipf’s law (also known as Z distribution) is a discrete variant of the Pareto distribution that in turn is a continuous power law.

Many between-sample normalization methods have been proposed for bulk and single-cell RNA-Seq data, and several attempts have been made to determine the best normalization procedure (Cole *et al*., 2019; Dillies *et al*., 2013; Evans *et al*., 2018; Tian *et al*., 2019). The general conclusion of these studies is that different datasets require different normalization strategies, and that the performance of normalization is influenced by many dataset-specific characteristics, such as sample heterogeneity, library preparation protocol, and sequencing depth.

Apart from the statistical aspects, single-cell RNA sequencing (scRNA-seq) has posed new considerable computational challenges. The increase in the number of cells per experiment translates into a dramatic increase in the data points to be analyzed, requiring methods able to efficiently scale to millions of cells, both in terms of memory usage and computational time. Typically, each step of the analysis, from normalization to clustering and functional analyses, can be highly demanding when dealing with hundreds of thousands or even millions of cells (Hicks *et al*., 2021; Lähnemann *et al*., 2020). In this perspective, a desirable normalization method should be able to scale efficiently with the number of cells, while simultaneously maintaining a good performance.

In the analysis of bulk and single-cell RNA-seq data, two major classes of between-sample normalization methods have been proposed: global scaling and non-linear approaches. The simplest scaling method is the Count Per Million (CPM) transformation, which simply scales the observed read (or UMI) counts by the total number of sequenced reads (or UMIs) per sample. More robust scaling procedures have been proposed in the bulk RNA-seq literature, such as TMM (Robinson and Oshlack, 2010), geometric mean scaling (DESeq2; Anders and Huber, 2010), and upper-quartile scaling (Bullard *et al*., 2010). In the context of single cell data, a popular scaling approach is the deconvolution strategy proposed in Lun *et al.* (2016a) and implemented in the *scran* Bioconductor package (Lun *et al*., 2016b). *Linnorm* (Yip *et al*., 2017), a linear model-based scaling algorithm, although not as popular as *scran*, has been shown to outperform other methods in a recent benchmark (Tian *et al*., 2019). More recently, *sctransform* (Hafemeister and Satija, 2019) *has gained popularity due to its good performance and its integration in the popular Seurat* package (Stuart *et al*., 2019). Briefly, *sctransform* uses the Pearson residuals of a regularized negative binomial model as normalized data.

While CPM is scalable to millions of cells, its performance is not always optimal (Robinson and Oshlack, 2010; Tian *et al*., 2019; Hafemeister and Satija, 2019); on the other hand, more robust normalizations, such as *scran* and *sctransform*, require a large amount of time and/or memory in big datasets.

Here we propose *PsiNorm*, a new scRNA-seq scaling normalization method, inspired by the Pareto power-law distribution. We compare *PsiNorm* to state-of-the-art methods in terms of concordance, scalability, and computational efficiency, as well as in terms of the accuracy of downstream clustering. We show that *PsiNorm* is the most scalable normalization among those that show good accuracy, being highly efficient in terms of memory usage and computational time. In particular, *PsiNorm* leads to comparable, and sometimes better, clustering than state-of-the art methods, such as *scran* and *Linnorm*, that either take longer or need more RAM. Finally, the ability of *PsiNorm* to work with out-of-memory data, such as HDF5 files, allows it to efficiently normalize datasets that may not even fit in RAM memory.

## 2 Approach and rationale

scRNA-seq data structures substantially differ from bulk. Potential gene dropouts and shallow sequencing make single cell data highly sparse. Moreover, the “large *p*, small *n*” paradigm (*p* being the number of genes, *n* the number of samples) that is typical of bulk data, is quickly moving towards the opposite scenario (*n > p*) with recent indexing-based experimental protocols. With the dramatic increase in the number of cells, all the analyses steps, including normalization, have become computationally intensive.

While some evidence showed a good fit of power-law distributions on bulk gene expression data, only few attempts have been made to fit such distributions to single-cell data (Townes and Irizarry, 2020). Motivated by these observations, here we investigate if and how power-law distributions could resemble scRNA-seq data empirical distributions with the goal of normalization in mind.

In the following i) we investigate the goodness-of-fit of the Pareto (type I) and Z (Zipf’s law) distributions on scRNA-seq data and ii) we propose a new method, called *PsiNorm*, to normalize raw read counts based on this fit. Then, iii) we compare our normalization in terms of cluster identification, concordance and computational resources required (time and memory usage) with other methods, proposed for bulk RNA-seq, such as logCPM, TMM and DESeq2, compositional data, such as the Centered Log Ratio (CLR), and scRNA-seq, such as Linnorm, sctransform and scran. The choice of these normalization methods represents a comprehensive set of methods that either have shown good performance in benchmark studies (e.g. Tian *et al*., 2019) or are popular among practitioners for the ease-of-use of their implementation.

## 3 Methods

### 3.1 The Pareto Distribution

The Pareto (type I) distribution is a continuous power-law probability distribution with support on the positive real axis. Its cumulative distribution function (cdf) is:

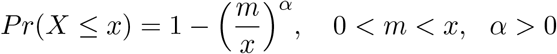

where *α* is the shape parameter and *m* is the minimum value of X.

The Pareto’s density function can be expressed as a power-law

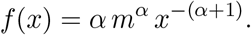

Given a sample of *n* independent observations, the parameter *α* can be estimated using the maximum likelihood method obtaining

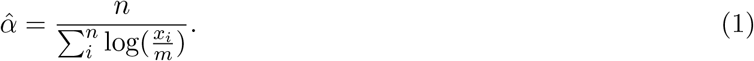

One important problem of fitting such distribution to sequencing data is that the Pareto distribution is defined only for *m >* 0, a condition not met since we always expect some genes with zero mapped reads.

There are two possible solutions to this problem. The first one (that we called Pareto0) estimates *α* on non-zero counts, while the second one (called Pareto+1) fits the model on pseudo-counts (raw counts +1). In this second approach 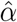 can be seen as the inverse of the log geometric mean of the pseudo-sample:

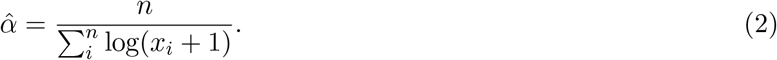

### 3.2 The Zipf’s law and its relation to Pareto

The Zipf’s power-law distribution originates from the observation that the frequencies of words in a text are inversely proportional to their ranks (Powers, 1998). It is a discrete distribution based on ranks and its probability mass function is given by:

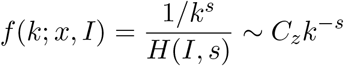

where *I* is the number of elements, *k* the vector of their ranks and *s* the coefficient characterizing the distribution. *H*(*I, s*) is the generalized harmonic series. Both Pareto and Zipf distributions are simple power laws with negative exponent and Zipf can be derived from the Pareto distribution if *X* values are binned into *I* ranks (Meintanis, 2009; Arnold, 2015).

Given the relationship between the two distributions, we can derive that *α* = 1*/s* (see Supplementary Text for details) (Meintanis, 2009; Arnold, 2015). However, while the maximum likelihood estimator of the Pareto *α* parameter has a closed-form, Zipf’s distribution parameter does not. Hence, numerical optimization methods are required.

### 3.3 The *PsiNorm* Normalization

The Pareto parameter *α* is inversely proportional to the sequencing depth, it is sample specific and its estimate can be obtained for each cell independently. Denoting by *X* the *I × J* matrix of read counts, with *I* genes and *J* cells, then the vector of normalized counts of cell *j*, 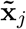, is equal to:

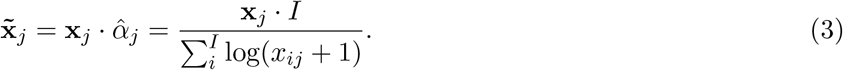

Given the inverse relationship between *α* and the sequencing depth, here 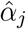 is used as a multiplicative normalization factor. We note that this essentially reduces to dividing each count by the sum of the log-counts of each cell, rescaled by a constant, a very similar approach to the CPM normalization. Note that often (e.g., in clustering and dimensionality reduction) it is useful to work with log-normalized counts. In the following, we will denote with log-normalized counts the quantity 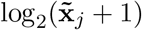.

In the following, *PsiNorm* is compared with seven state-of-the-art methods (see Supplementary Text), in terms of clustering performance, concordance, and computational efficiency.

### 3.4 Evaluation Criteria

#### 3.4.1 Cluster analyses

To evaluate the ability of normalization to remove technical bias and reveal the true cell similarity structure, we used both an unsupervised and a supervised approach, since we know the labels of the datasets used for the comparison (see Section 3.5 for details).

In the unsupervised approach, we applied principal component analysis (PCA) on the log-normalized counts and, using the first 50 PCs, we identified clusters using a partitional method (*clara* in the *cluster* R package) with *k* (number of groups) equal to the known number of clusters. Then, we computed the Adjusted Rand Index (ARI) to compare the known and the estimated partition (Hubert and Arabie, 1985).

In the supervised approach, we computed the silhouette index of the known partition in the reduced dimensional space obtained by PCA of the log-normalized counts. The rational is that a normalization that properly reduces technical noise should lead to compact clusters with high cohesion and separation that correspond to the known cell populations.

#### 3.4.2 Concordance Analyses

We estimated within-method concordance (replicability) by randomly splitting each dataset into two equally sized parts and evaluating the cardinality of the intersection between the two lists of most variable genes after normalization. The splitting is repeated 10 times and the average within-method concordance is reported. Between-dataset concordance (reproducibility) has been evaluated using scRNA-seq data of the same samples obtained with different experimental techniques. The cardinality of the intersection between the two lists of most variable genes after normalization across dataset is used as a measure of concordance.

### 3.5 Real Datasets

We used two sets of data to compare methods: the scRNA-seq mixed human cell lines experiments from Tian *et al.* (2019), which we refer to as the *mixology* dataset, and the mouse primary motor cortex datasets generated by the BRAIN Initiative Cell Census Network (BICCN) (Yao *et al*., 2020), which we refer to as the *BICCN* dataset.

In the mixology dataset, five human lung adenocarcinoma cell lines were cultured separately, single cells from each cell line were mixed in equal proportions, with libraries generated using three different protocols CEL-seq2, Drop-seq with Dolomite equipment and 10X Chromium (Tian *et al*., 2019).

In the BICCN dataset, over 700,000 cells were characterized via single-cell and single-nucleus RNA-seq (using 10X and SMART-seq protocols) to comprehensively identify all cell types in the adult mouse primary motor cortex (Yao *et al*., 2020).

To compare the normalization approaches in terms of concordance and clustering performance, we selected a random subset of 500 cells from both single-cell and single-nucleus samples for each sequencing protocols 10X v2, 10X v3 and SMART-Seq. All datasets were filtered to keep only those genes with more than 2 reads in more than 5% of cells and discarding cells without labels. For further details see Supplementary Table S1.

### 3.6 Case Study

As a case study, we use the complete 10X v2 BICCN dataset. After a filtering procedure that retained the genes with more than 2 reads in more than 5% of cells and discarded the cells without labels, we obtained a matrix with 7,171 genes and 124,330 cells. Cell labels were provided by Yao *et al.* (2020) with three different degree of details: *cluster* (a fine-grained partition that contains cell sub-populations), *sub-class* (which defines the major cell types of the adult mouse motor cortex), and *class* (which partitions the cells in broad classes, i.e., excitatory neurons, inhibitory neurons, and non-neuronal cells). We used the *sub-class* label as our ground truth.

Clusters were identified as in Section 3.4.1. Then, we used the ARI to compare the cluster identified after each normalization with the known labels.

### 3.7 Simulated Datasets

We used simulations to compare normalization methods in terms of their computational efficiency (RAM usage and CPU time). To simulate data we used the *splatSimulateSingle* function of the *splatter* R/Bioconductor package (Zappia *et al*., 2017), with default parameters. We set the number of genes equal to 10,000 with increasing number of cells: 25,000, 50,000, 75,000 and 100,000 cells. RAM usage and computational time were recorded for a single core usage.

### 3.8 Software and data availability

An implementation of *PsiNorm* is available in the R package *PsiNorm* available at https://github.com/MatteoBlla/PsiNorm. The code to generate the analysis and figures of this manuscript is available at https://github.com/MatteoBlla/PsiNorm-plot.

The mixology dataset is available at https://github.com/LuyiTian/sc_mixology. The BICCN dataset was generated as part of the BICCN consortium and can be downloaded from http://data.nemoarchive.org/biccn/lab/zeng/transcriptome/.

## 4 Results

### 4.1 Goodness-of-fit

To evaluate the goodness of fit of the Pareto and Zipf models on single cell data we evaluated two distinct aspects: i) the power-law fit and ii) the differences between expected and observed counts. We first visually inspected the log-log plot of expression versus rank to check the approximation to a power law for three classes of cells characterized by the minimum, median and maximum sequencing depths and for different technologies (Fig. 1A). Secondly, for each cell, we estimated the parameters of Pareto0, Pareto+1 and Zipf distributions. Using these estimates, we compared the log ratio between the theoretical and the empirical third quartiles; the closer this ratio is to 0, the better the goodness of fit (Fig. 1B).

**Figure 1:**
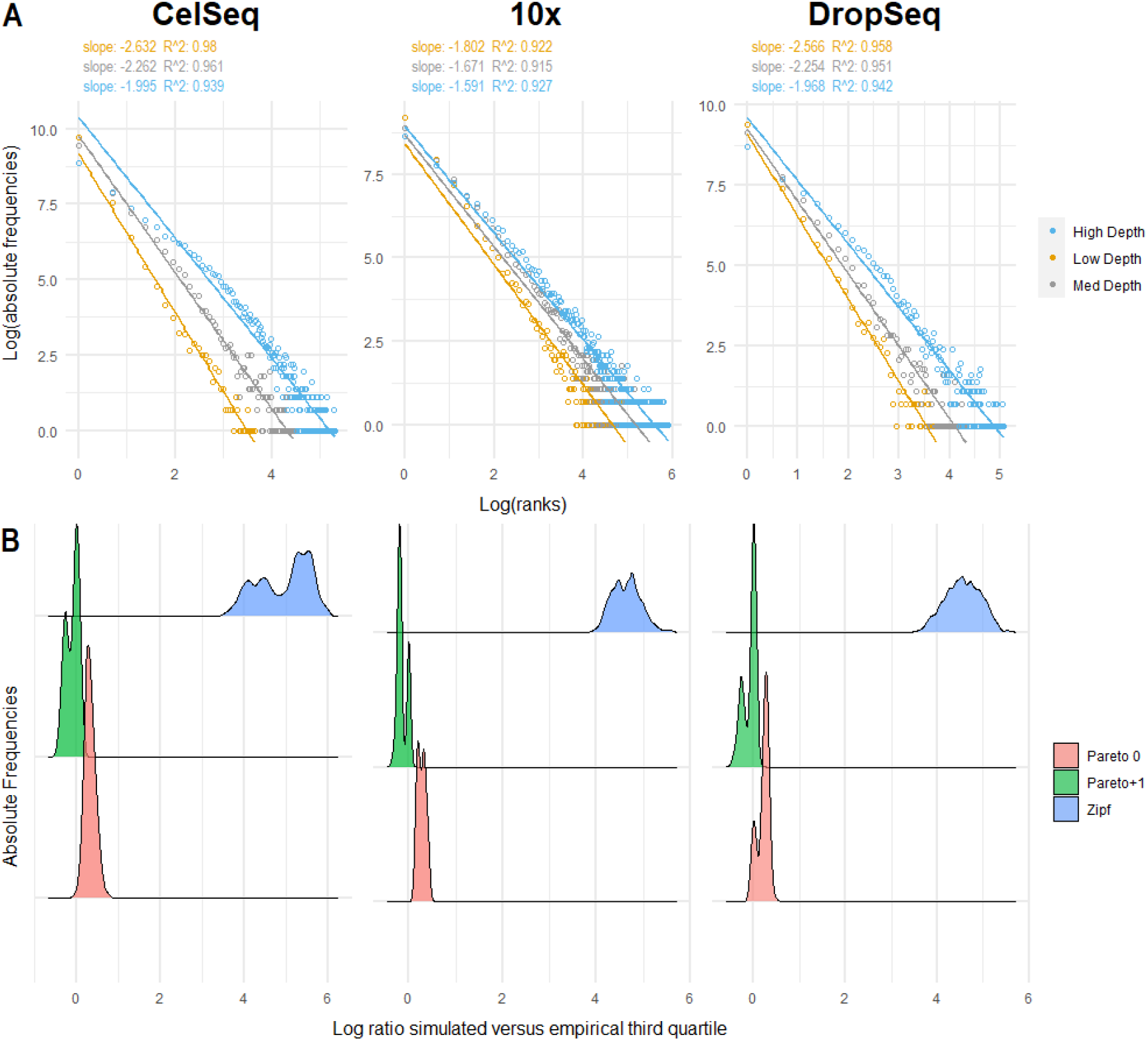
Goodness of fit. **Panel A**. Log-expression vs log-frequency plot of the cells with the minimum, median and maximum depth per technology. Linear fit is reported. **Panel B**. Distribution of the log ratios between simulated and empirical third quartiles per cell across different technologies.

Single-cell data are well approximated by a power-law (*R*^2^ *>* 0.9) independently of the sequencing depths and technology (Fig. 1A and Supplementary Fig. 1A). However while Zipf’s law largely overestimates counts, the Pareto distribution is more flexible and better fits scRNA-seq data, as shown by the distribution of the log ratios of the simulated versus empirical third quartiles (Fig. 1B and Supplementary Fig. 1B). Indeed, while Zipf’s law over-estimates the third quartile of the distribution, the log ratio between Pareto simulated and empirical third quartiles is close to zero (Fig. 1B). In particular, the use of the pseudo-counts for parameter estimation (Pareto+1) shows a better goodness-of-fit than the removal of zero counts (Pareto0; Fig. 1B).

Taken together our results indicate that the Pareto distribution on pseudo-counts well resembles scRNA-seq data, independently of sequencing depths and technology.

### 4.2 *PsiNorm* leads to comparable distributions across cells

Figure2A shows the effect that *PsiNorm* has on the expression distribution on three representative cells (with low, moderate, and high depths). After normalization, the distributions of the highly expressed genes (those with small ranks) are aligned. The effect on the entire dataset can be appreciated in Figure2B where the slope and intercept distributions are reported for raw and normalized data. As expected, after normalization the variability of both distributions is greatly reduced.

**Figure 2:**
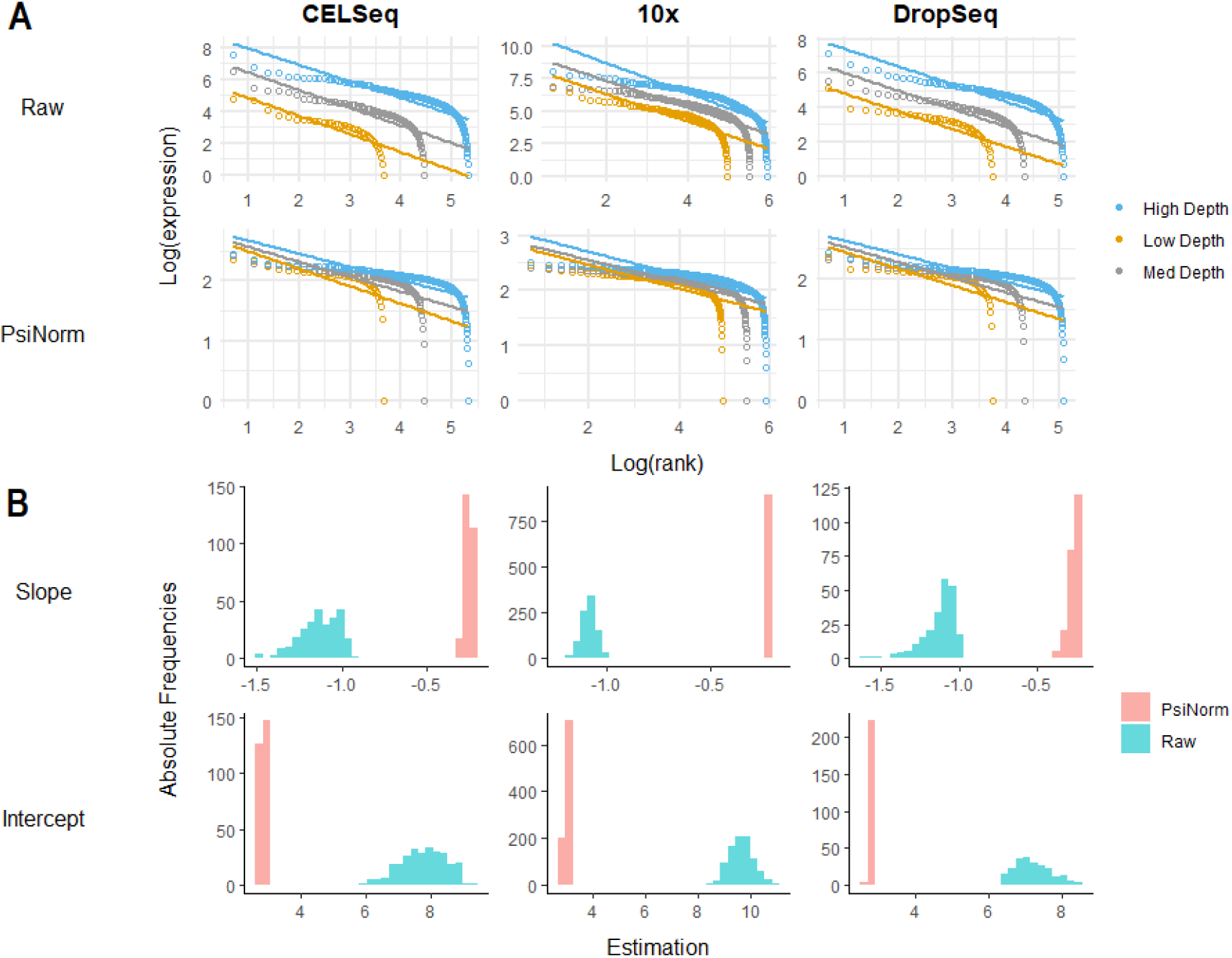
Pareto normalization. **Panel A**. The log expression ordered from the highest to the lowest of three classes of cells (with low, moderate and high coverage) is reported for raw and Pareto normalized data. The linear fit is reported for each cell. **Panel B**. The density distributions (across all cells per technology) of the linear fit estimates (slopes and intercepts) of raw and normalized data.

These findings confirm that *PsiNorm* is able to effectively scale the data making the distribution of highly expressed genes comparable across cells.

### 4.3 Impact of normalization on cell clustering

Organizing cells into groups is the first intermediate result of any single-cell analysis. Here we wonder whether *PsiNorm* transforms the data maintaining the similarity structure among cells, allowing a downstream clustering algorithm to detect cell populations.

Figure 3A shows an example of principal component analysis (PCA) obtained with different normalizations in the mixology dataset with five groups (CELSeq2 5cl p3). See Supplementary Fig S3-S8 for the other datasets.

**Figure 3:**
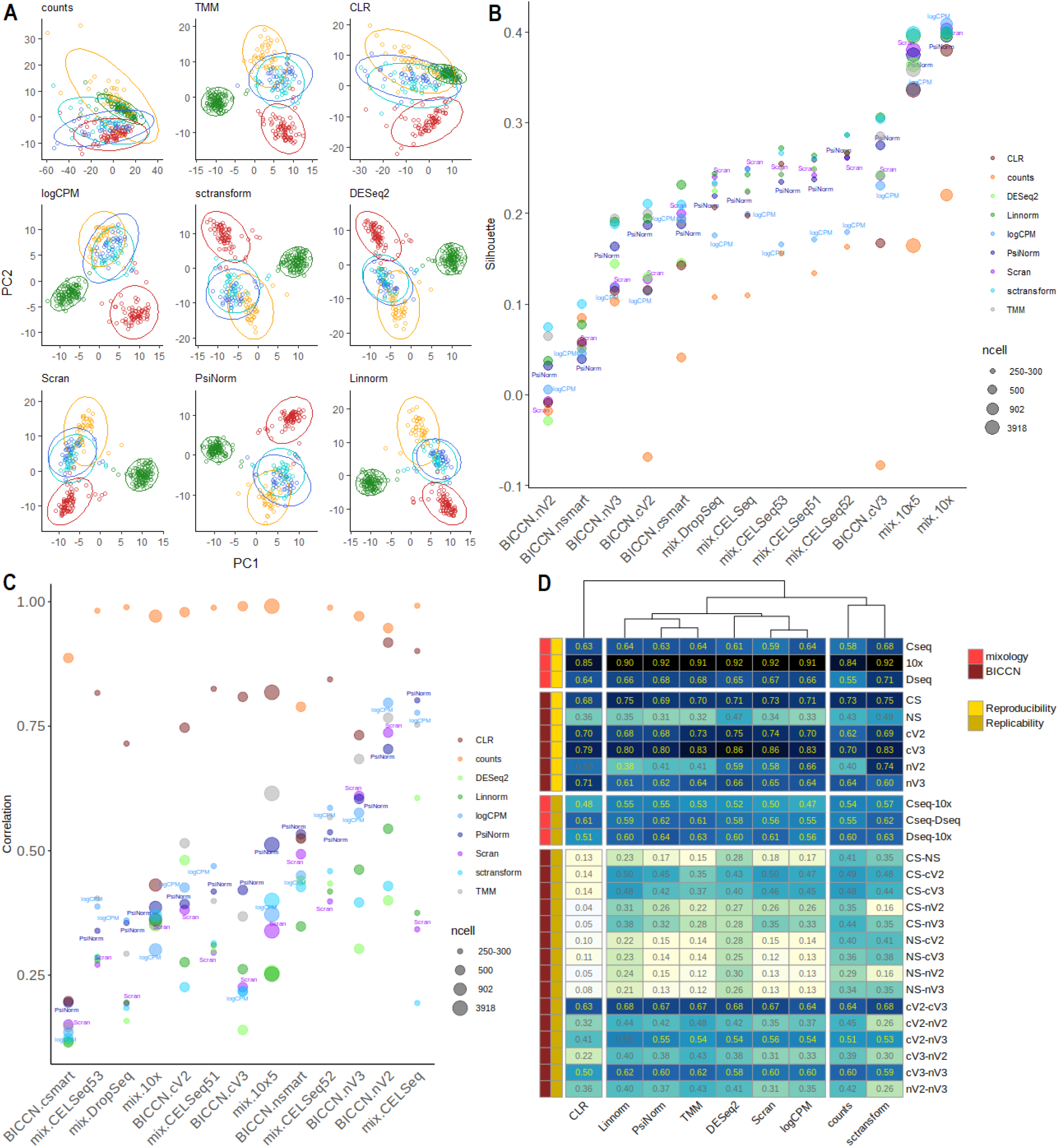
**Panel A**. Principal component analyses (PC1 vs PC2) of CELSeq3 dataset composed of 5 groups (highlited with different colours). See Supplementary Figure S3-S8 for PCA analyses on the other datasets. **Panel B**. Silhouette index across different dataset and different normalization methods. Datasets are sorted by the silhouette index obtained with the Pareto normalized data. The dot dimension is proportional to the dimension of the datasets in terms of number of cells. **Panel C**. The maximum correlation index between PC1 and PC2 and cell sequencing depths is reported for each dataset **Panel D**. The upper six raws of the heatmap shows the degree of replicability in colour scale, the number within cells is the average concordance between the top variable genes in ten-fold random splits of the dataset. The lower fifteen raws show reproducibility in colour scale, the number within cells is the concordance between the top variable genes in different datasets of the same sample but obtained with different technology.

Apart from CLR that hardly recognize the similarity structure of the known groups, all the other methods are able to identify the major differences among the cell lines (Fig 3A). This is confirmed in all other datasets. Interestingly, logCPM and sctransform did not perform well in the 5 class 10X dataset (Supplementary Fig. 4A).

We computed the ARI of all partitions to compare the inferred clusters and the real cell line classification. Linnorm and sctransform lead to the highest ARI, followed by *PsiNorm* and scran (Table 1).

**Table 1:** Normalization evaluation. Computational costs and RAM usage are referred to the simulation matrix with 100,000 cells.

|  | average<br>ARI | average<br>silhouette | average correlation<br>PCA-depth | average within<br>concordance | average between<br>concordance | computational<br>costs (sec) | RAM usage<br>(Gb) |
| --- | --- | --- | --- | --- | --- | --- | --- |
| sctransform | 0.756 | **0.244** | **0.310** | **0.712** | 0.417 | 1154 | 43 |
| Linnorm | **0.767** | 0.241 | 0.323 | 0.641 | **0.424** | 446 | 23 |
| PsiNorm | 0.734 | 0.218 | 0.476 | 0.638 | 0.388 | 112 | 17 |
| Scran | 0.720 | 0.212 | 0.369 | 0.674 | 0.375 | 1838 | 16 |
| TMM | 0.711 | 0.228 | 0.491 | 0.651 | 0.378 | 506 | 25 |
| logCPM | 0.696 | 0.180 | 0.450 | 0.676 | 0.368 | **64** | **12** |
| DESeq2 | 0.695 | 0.204 | 0.329 | 0.692 | 0.414 | 192 | 20 |
| CLR | 0.626 | 0.191 | 0.714 | 0.661 | 0.270 | 122 | 40 |

Exploiting known cell labels, we used the silhouette width to quantify the cohesion of the clusters and the separation of the cell lines. Figure 3B shows the average silhouette widths for each normalization-dataset combination. In general, single nucleus datasets show a lower average silhouette, independently of the normalization (Fig. 3B). This is probably due to the higher level of sparsity that characterize these data. Furthermore, the average silhouette depends on the number of cells (the more cells the higher the silhouette) and, perhaps unsurprisingly, on the complexity of the dataset: the simple mix of cell lines from the mixology dataset showed a higher silhouette than the complex BICCN data (Fig. 3B). In terms of normalization performance, our analysis confirmed that no single method outperforms all others in all datasets: for instance scran, which was among the top performers in the mixology 10X datasets, did not perform as well in the BICCN 10X v2 datasets (both single-cell and single-nucleus). Overall, Linnorm, sctransform, TMM and *PsiNorm* showed the most consistent performance (Table 1).

When a normalization fails to reduce unwanted variation within a dataset (due for instance to differences in sequencing depth), the factors computed by the dimension reduction technique might capture technical noise rather than biological variability. To check whether the first two PCs are capturing technical variance, we computed the maximum correlation obtained between PC1 and PC2 and cell sequencing depths (Fig. 3C). A higher correlation indicates that the normalization was not able to properly remove noise.

While we observed a general high correlation for CLR (and no normalization), TMM shows high correlations only for some datasets confirming that these methods do not remove enough technical variation (Fig. 3C). All other methods performed similarly, with sctransform, DESeq2 and Linnorm as top performers (Fig. 3C and Table 1).

### 4.4 Concordance Analyses

Replicability and reproducibility are two important aspects when dealing with data transformations. Here we defined replicability as the ability to maintain the order of the most variable genes between two random split of the same dataset (within-dataset concordance) and reproducibility as the ability to maintain the order of the most variable genes between two independent datasets measuring the same samples (between-dataset concordance).

As expected, we observed a general higher concordance within than between datasets (Fig. 3D). Indeed, the mean of the within-dataset concordance was 0.72 for the mixology dataset and 0.62 for the BICCN data. On the other hand, the average between-dataset concordance was 0.57 for the mixology dataset and 0.34 for the BICCN data. Single-nucleus datasets showed the lowest within-dataset concordance while the 10X mixology dataset showed the highest (Fig. 3D). We observed similar results for the between-dataset concordance. As expected, between-dataset concordance was higher for datasets from similar platform, e.g., 10X and Dropseq showed a higher concordance than 10X and SMART-seq (Fig. 3D). Interestingly, the concordance between single-cell 10X V2 and V3 was higher than that between single-cell 10X V2 and single-nucleus 10X V2 (and between single-cell 10X V3 and single-nucleus 10X V3), suggesting that the 10X chemistry was less important than the RNA provenance in determining concordance (Fig. 3D).

In terms of normalization performance, methods fell into three main groups: CLR showed lower level of both within- and between-dataset concordance; raw counts and sctransform showed a high between-dataset concordance; and *PsiNorm*, Linnorm, TMM, Scran, DESeq2 and logCPM performed well both in term of within- and between-dataset concordance (Fig. 3D and Table 1).

### 4.5 Computational performance

To complete our benchmark, we performed a comparative analyses of computational performances in different simulated settings. Figure 4 and Table1 show the RAM usage and the elapsed time for each method. As expected, logCPM, PsiNorm, and DESeq2 are the most scalable methods (Fig. 4). Indeed these methods only need simple operations (such as averages and multiplications) to scale the data. While CLR is as fast as the above-mentioned methods, it is much more demanding in terms of memory usage. At the other end of the spectrum, scran and sctransform are not as scalable. sctransfrom requires the highest amount of RAM among the tested methods, exceeding 40 GB for 100,000 cells (Fig. 4). While scran is much more memory-efficient, it is the slowest method, requiring about 30 minutes for 100,000 cells (Fig. 4). Overall, *PsiNorm* is very scalable, being only slightly slower than the simplest strategy (logCPM).

**Figure 4:**
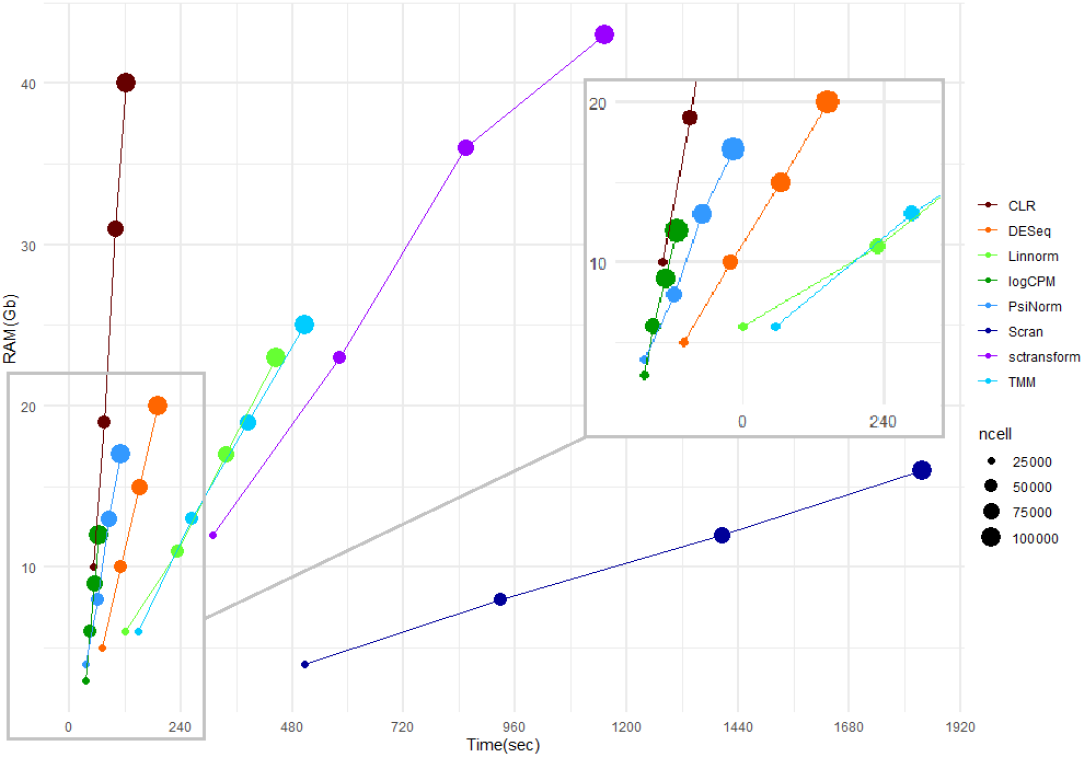
Computational performance. Comparative evaluation of RAM usage and computational time on simulated data with increasing number of cell.

### 4.6 Case Study

As a case study, we analyzed the full 10X V2 single-cell data from the BICCN study (Yao *et al*., 2020). These data consists of 124,330 cells and 7,171 genes.

We applied the four methods that showed the best performance among those with a limited memory footprint, i.e., Linnorm, scran, logCPM, and *PsiNorm*. Although sctransform performed well in our benchmark, its memory usage prevents its use in very large datasets.

Figure 5 shows the UMAP plot obtained after each normalization. While all methods are able to separate the major cell types, the comparison with the BICCN labels showed that scran and *PsiNorm* lead to the best agreement in terms of ARI (Fig. 5). Scran is confirmed to be the most time consuming method, taking more than 30 minutes to normalize the matrix. On the other hand, *PsiNorm* is almost as fast as logCPM, completing the task in just under 3 minutes.

**Figure 5:**
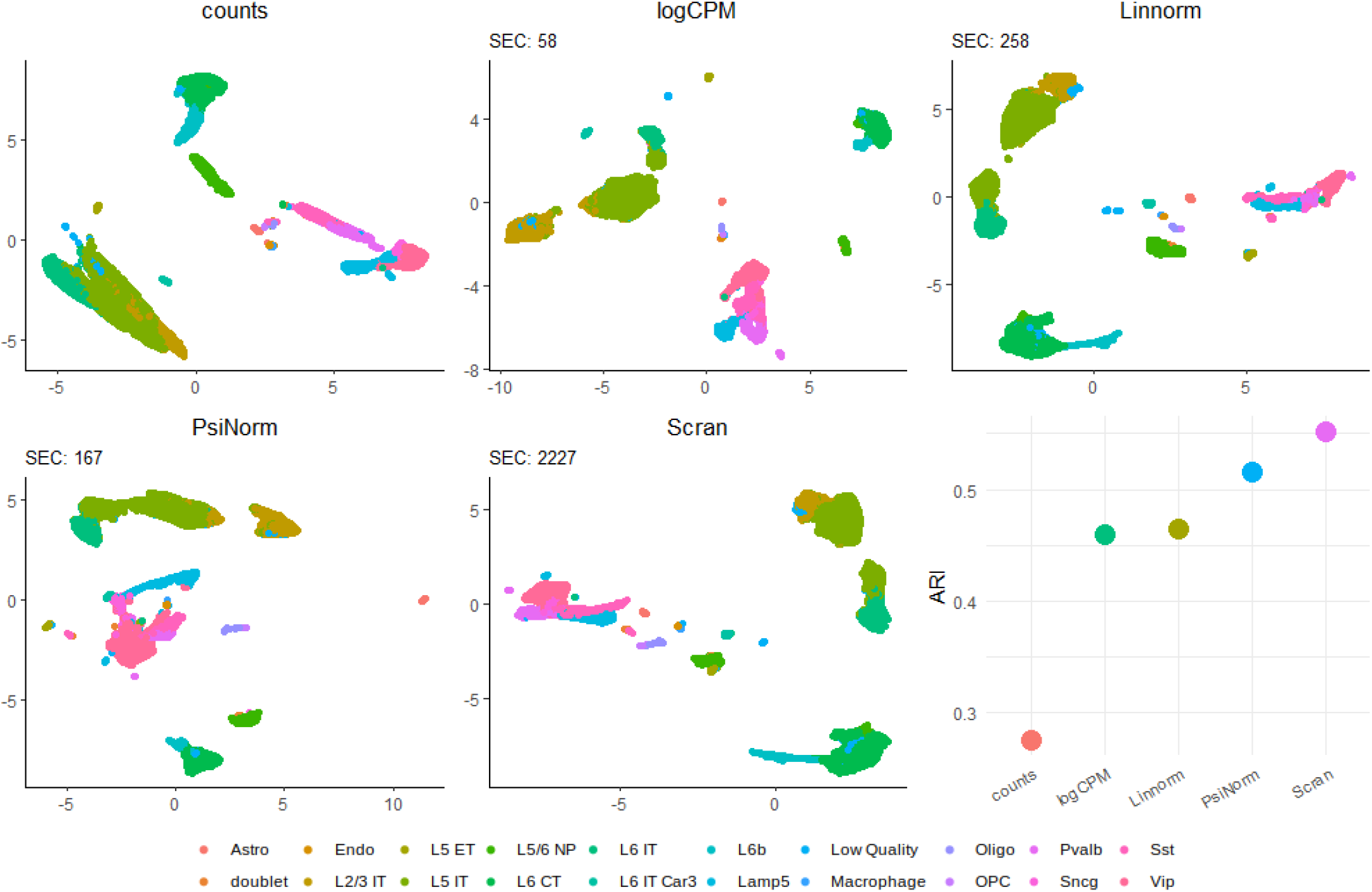
UMAP plots of the case study data obtained with raw data and after the four best performing normalizations. The Adjusted Rand Index comparing the inferred vs the known groups are reported.

## 5 Discussion

In single-cell experiments, computational efficiency in term of time and memory usage is a key aspect. The massive number of cells, combined with the large number of genes make even simple scaling normalization demanding. For instance, scran applied to a dataset of 1.3 million datasets take more than 5 hours (Hicks *et al*., 2021).

Based on the Pareto distribution scale parameter estimate, 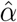, we derived a simple and scalable global between-sample normalization method, called *PsiNorm. PsiNorm* is fast and memory efficient. Moreover, through the integration with the Bioconductor *DelayedArray* framework (Pagès *et al*., 2019), it can be applied to dense or sparse in-memory matrices as well as out-of-memory data representations, such as data stored in HDF5 files (The HDF Group, 1997).

*PsiNorm* does not need a reference and is performed independently for each cell. This is useful for supervised classification settings, in which it can be useful to apply normalization to new out-of-sample data. The final goal of the transformation is to align the gene expression distribution especially for those genes characterised by high expression. Note that, similar to other global scaling methods, our method does not remove batch effects, which can be dealt with downstream tools (e.g., Risso *et al*., 2014; Haghverdi *et al*., 2018; Butler *et al*., 2018).

Globally our results are summarized in Table 1, where the best method for each task is reported in bold. We observed that, as expected, normalizations specifically designed for scRNA-seq data are among the best performing. Among them we found the *PsiNorm* and scran show good performances in six features.

To conclude, normalization for the purpose of clustering and cell type discovery seems less critical than normalization for differential expression, and even very simple methods, such as logCPM, work well in several cases. Hence, methods’ scalability becomes an important aspect to consider in the choice of normalization. Our proposed *PsiNorm* normalization showed a good trade-off between accuracy and scalability, making it a promising method for very large datasets.

## Acknowledgements

We thank Hongkui Zeng and the members of the BICCN consortium for sharing the BICCN dataset.

