## Supplementary material for "PsiNorm: a scalable normalization for single-cell RNA-seq data"

April 7, 2021

#### 1 The Zipf's law and its relation to Pareto

The Zipf's power-law is a power-law discrete distribution based on ranks whose probability mass function is given by:

$$f(k; x, I) = \frac{1/k^s}{H(I, s)} \sim C_z k^{-s}$$

where $I$ is the number of elements, $k$ the vector of their ranks and $s$ the coefficient characterizing the distribution. $H(I, s)$ is the generalized harmonic series.

The estimation of $s$ is not available in closed form and it is necessary to estimate it through recursive optimization methods. We used the *stats4::mle* R function to compute the estimate of $s$ as the value that maximizes the likelihood function. We used Nelder and Mead (1965) optimization method (the default of *optim* function) which uses the concept of simplex to approximates a local optimum.

Both Pareto and Zipf distributions are simple power laws with negative exponent and Zipf can be derived from the Pareto distribution if $X$ is a Pareto random variables and its values are binned into $I$ ranks.

Specifically, the Pareto's density function $f(x)$ is a power-law:

$$f(x) = \alpha m^{\alpha} x^{-(\alpha+1)} = C_p x^{-\beta}$$

where $C_p = \alpha m^{\alpha}$ and the parameter of a generic power-law $\beta$ is equal to $\alpha + 1$.

Then, the mean of the $k$-th random variable $X$ distributed as Zipf is equal to $E[X_k] \sim C_1 \times k^{-s}$ with the meaning that there are $k$ variables with the expected value higher than this quantity. We obtain that $Pr[X \geq C_1 \times k^{-s}] = C_2 \times k$ and given $y = k^{-s}$ and deriving the distribution function the result is $P[X = y] \sim y^{-1+(1/s)} = y^{-\beta}$. So from the relationships between $\alpha$ and $\beta$ and between $\beta$ and $s$ we obtain:

$$\beta = \alpha + 1 = 1 + 1/s \Rightarrow \alpha = 1/s$$

[1] Department of Biology, University of Padova, Italy.
[2] Department of Statistical Sciences, University of Padova, Italy.

#### 2 Banchmarked methods

**Count per million (CPM).** This method simply divides read counts by the sequencing depth defined as the sum of the expression of the genes per cell. Each count is then multiplied by a million to make normalized count not too much compressed. Usually, the base 2 logarithm of the normalized pseudo-count is taken, defining the *logCPM* values:

$$\tilde{x}_{ij} = log_2\Big(\frac{x_{ij} \times 10^6}{N_j} + 1\Big)$$

with $N_j = \sum_i x_{ij}$ the sum of the counts of cell $j$.

**Centered Log-Ratio (CLR).** CLR is similar to logCPM with the difference that it divides pseudo-counts by the geometric mean of each cell. Given $gm_{x_j+1}$ the geometric mean of the $j$-th cell:

$$gm_{x_j+1} = \Big(\prod_{i=1}^{n}(x_{ij}+1)\Big)^{\frac{1}{n}}$$

the normalized counts are:

$$\tilde{x}_{ij} = log\Big(\frac{x_{ij}}{gm_{x_j+1}} + 1\Big)$$

**scran.** scran is based on a cell pooling strategy. Given the global reference $\overline{x}$ defined as:

$$\overline{x} = \frac{1}{J}\sum_j x_{ij}, \quad i = 1, ..., n, \quad j = 1, ..., J$$

and $k$ overlapping groups of cells, scran estimates the size factor $SF_{p_k}$ of each pool under the assumption that every $SF_{p_k}$ is a linear combination of the size-factors of the cells that belong to the pool:

$$\forall\ pool_k \ : \sum_{j \in p_k} x_{ij} = \Big[x_{1p_k}, ..., x_{np_k}\Big]$$

$$SF_{p_k} = Median\Big(\frac{x_{1p_k}}{\overline{x}_1}, ..., \frac{x_{np_k}}{\overline{x}_n}\Big) = \sum_{j \in p_k} SF_j$$

Solving the equations, we obtain size factors for each cell and define the normalized values as:

$$\tilde{x}_{ij} = log\Big(\frac{x_{ij}}{SF_j} + 1\Big)$$

**DESeq2** Deseq2 uses as reference the geometric means of the of gene across cells. For every gene $i$:

$$gm_{x_i} = \Big(\prod_{j=1}^{J} x_{ij}\Big)^{\frac{1}{J}}$$

Then each count is divided by its geometric mean and the median of these ratios is the size factor for the sample $j$.

$$SF_j = Median\Big(\frac{x_{1j}}{gm_{x_1}}, ..., \frac{x_{nj}}{gm_{x_n}}\Big)$$

The normalized counts are obtained by taking the log of the ratio of each counts and its size factor:

$$\tilde{x}_{ij} = log_2\Big(\frac{x_{ij}}{SF_j} + 1\Big)$$

**Trimmed Mean of M-values (TMM).** TMM (Robinson and Oshlack, 2010) defines the log-fold-changes (M) and absolute expression levels (A) between each cell and a reference (by default the cell whose upper quartile is closest to the mean upper quartiles across cells):

$$M_{ij}^{(r)} = log_2\Big(\frac{x_{ij}/N_j}{x_{ir}/N_r}\Big)$$
$$A_{ij}^{(r)} = \frac{1}{2}log_2(x_{ij}/N_j * x_{ir}/N_r)$$

to apply a trimming procedure. By default the method trims the 30% of highest and lowest values for $M_{ij}$ and 5% of highest and lowest values for $A_{ij}$. After the trimming, the mean of $M_{ij}$ weighted by the inverse of the approximate asymptotic variances is used to normalize the counts:

$$log_2(SF_j^{(r)}) = \frac{\sum_{i \in I^*} M_{ij}^{(r)} w_{ij}^{(r)}}{\sum_{i \in I^*} w_{ij}^{(r)}}$$
$$w_{ij}^{(r)} = \frac{N_j - x_{ij}}{N_j x_{ij}} + \frac{N_r - x_{ir}}{N_r x_{ir}}$$
$$\tilde{x}_{ij} = log_2\Big(\frac{x_{ij}}{SF_j N_j} + 1\Big)$$

where $I^*$ is the set of genes with valid $M_{ij}$ and $A_{ij}$ values.

**Linnorm. Linnorm** (Yip *et al.*, 2017) filter genes according to their sparseness, variability and skewness in order to identify a set of stable genes. Then, given $R_{ij} = \frac{x_{ij}}{N_j}$ it defines the log of the normalized pseudo-counts as follow:

$$T_{ij} = ln(\lambda R_{ij} + 1)$$

The purpose is to identify the $\lambda$ (dataset-specific) that minimize the deviation from homoscedasticity and normality:

$$F(\lambda) = V(\lambda)^2 + S(\lambda)^2$$
$$\lambda = argmin(F(\lambda))$$

where $V(\lambda)$ represents the deviation of $T_{ij}$ from homoscedasticity and $S(\lambda)$ the deviation from the skewness of the dataset. Once $\hat{\lambda}$ has been obtained, *Linnorm* uses the quantities

$G_{ij} = ln(\hat{\lambda} R_{ij})$ to define $n$ regression models, $g_i = m_j x_{ij} + c_j$, where $g_i$ is the mean expression and $x_{ij}$ the sample's expression. Model parameters, $m$ and $c$, are updated with the equations $m^{updated} = \mu(m-1)+1$ and $c^{updated} = c \times \mu$ with $\mu$ set by default to 0.5 which provides a moderate level of normalization strength. Finally, given $B_{ij} = exp(m_j^{updated} G_{ij} + c_j^{updated})$ the counts are normalized:

$$\tilde{x}_{ij} = ln(B_{ij} + 1)$$

**sctransform.** **sctransform** is based on a regression model per gene with negative binomial error distribution and logarithmic link function ((Hafemeister and Satija, 2019)). For a given cell $j$ and gene $i$ it can define the expected counts and the expected standard deviation as follow:

$$log(\mu_{ij}) = \beta_{0_i} + \beta_{1_i} log_{10} N_j$$

$$\sigma_{ij} = \sqrt{\mu_{ij} + \frac{\mu_{ij}^2}{\theta_i}}$$

where $\beta_{0_i}$ and $\beta_{1_i}$ and the dispersion parameter $\theta$ have to be estimated. To avoid overfitting, SCT exploits the trend of the estimates versus gene mean to perform independent regularizations for all parameters. The regularized parameters are used to define the normalized counts as the Pearson residuals of the model:

$$\tilde{x}_{ij} = \frac{x_{ij} - \mu ij}{\sigma_{ij}}$$

where $\mu_{ij}$ is the expected count of gene i in cell j in the regularized negative binomial regression model, and $\sigma_{ij}$ is the expected standard deviation.

### 3 Supplementary Tables

Supplementary Table 1: Description of the datasets used to compare and evaluate PsiNorm normalization performances.

| Dataset name | N. genes | N. cells | N. clusters | % nulls | Technology | Sample type | Reference |
|---|---|---|---|---|---|---|---|
| 10x | 16468 | 902 | 3 | .45 | 10x | Cell mixture | mixology Tian et al. (2019) |
| CELSeq | 19759 | 274 | 3 | .64 | celseq | Cell mixture | mixology Tian et al. (2019) |
| DropSeq | 14947 | 225 | 3 | .62 | dropseq | Cell mixture | mixology Tian et al. (2019) |
| CELSeq51 | 15564 | 297 | 5 | .61 | celseq | Cell mixture | mixology Tian et al. (2019) |
| CELSeq52 | 14078 | 307 | 5 | .60 | celseq | Cell mixture | mixology Tian et al. (2019) |
| CELSeq53 | 13426 | 305 | 5 | .64 | celseq | Cell mixture | mixology Tian et al. (2019) |
| 10x5 | 11786 | 3918 | 5 | .63 | 10x | Cell mixture | mixology Tian et al. (2019) |
| csmart | 17998 | 500 | 14 | .53 | smart | cells | BICCN Zeng Yao et al (2020) |
| nsmart | 17902 | 500 | 17 | .73 | smart | nucleus | BICCN Zeng Yao et al (2020) |
| cV2 | 15784 | 500 | 17 | .73 | 10x | cells | BICCN Zeng Yao et al (2020) |
| cV3 | 16837 | 500 | 17 | .61 | 10x | cells | BICCN Zeng Yao et al (2020) |
| nV2 | 14791 | 500 | 14 | .89 | 10x | nucleus | BICCN Zeng Yao et al (2020) |
| nV3 | 15889 | 500 | 15 | .80 | 10x | nucleus | BICCN Zeng Yao et al (2020) |

### 4 Supplementary Figures

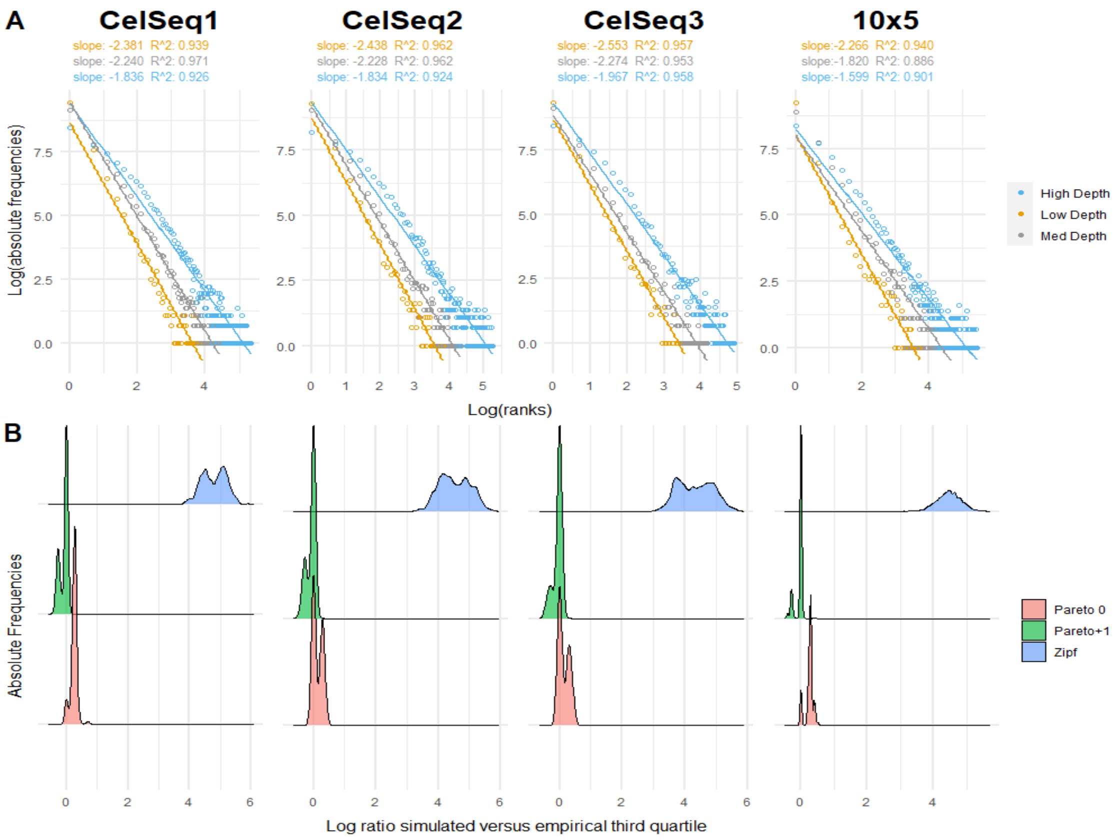


Supplementary Figure 1: Goodness of fit. **Panel A.** Log-expression vs log-frequency plot of the cells with the minimum, median and maximum depth per technology. Linear fit is reported. **Panel B.** Distribution of the log ratios between observed and expected third quartiles per cell across different technologies.

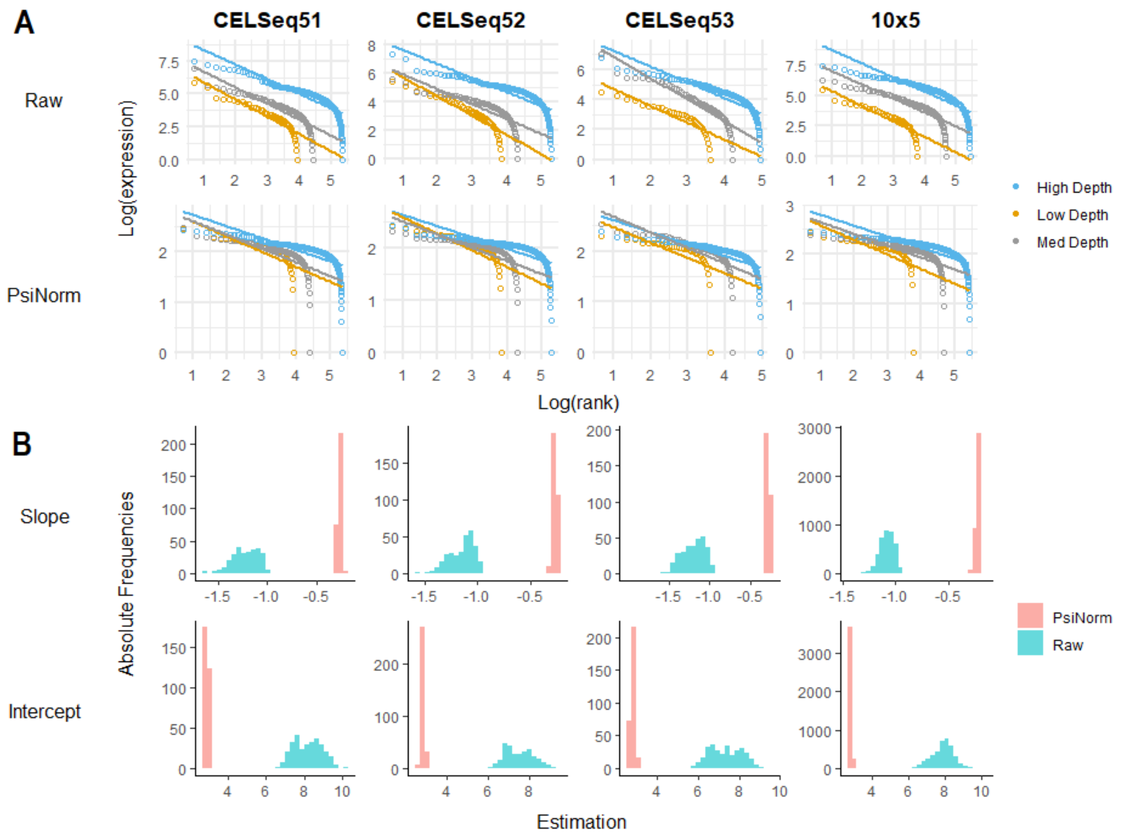


Supplementary Figure 2: Pareto normalization. **Panel A.** The log expression ordered from the highest to the lowest of three classes of cells (with low, moderate and high coverage) is reported for raw and Pareto normalized data. The linear fit is reported for each cell. **Panel B.** The density distributions (across all cells per technology) of the linear fit estimates (slopes and intercepts) of raw and normalized data.

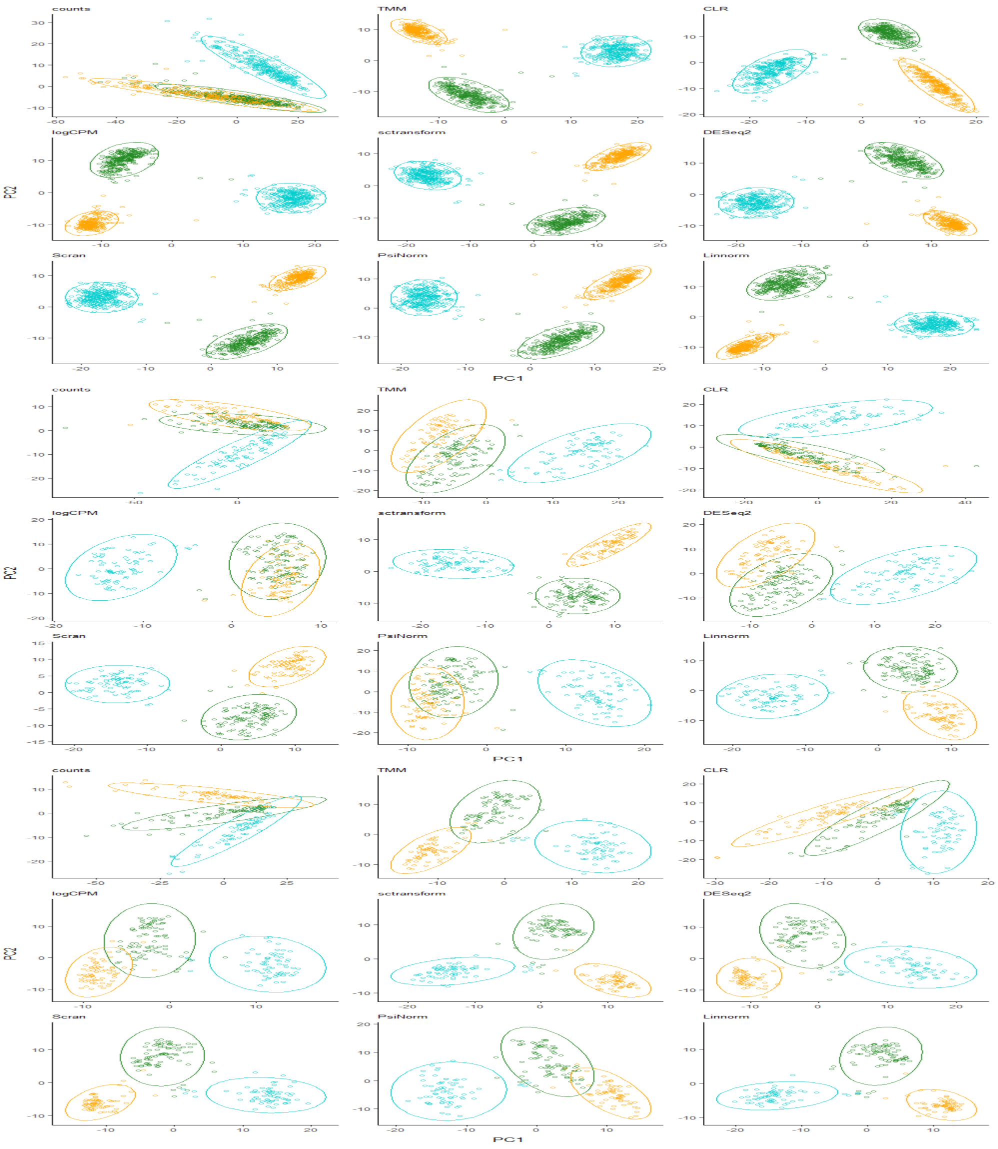


Supplementary Figure 3: PCA plot of the data with 3 cell types. **Panel A.** 10x data. **Panel B.** CELSeq data. **Panel C.** DropSeq data.

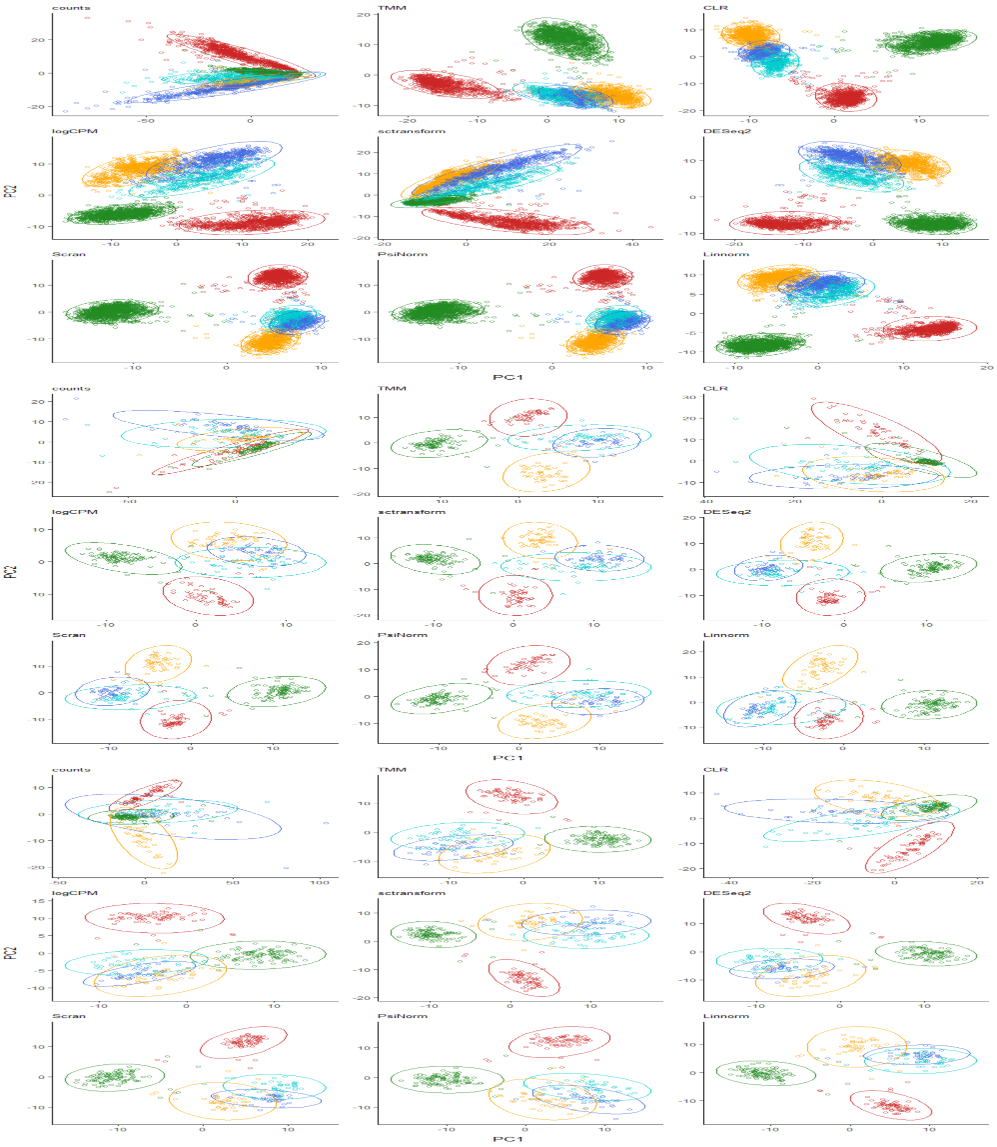


Supplementary Figure 4: PCA plot of the data with 5 cell types. **Panel A.** 10x5 data. **Panel B.** CELSeq1 data. **Panel C.** CELSeq2 data.
